# CRISPRi-Driven Osteogenesis in Adipose-Derived Stem Cells for Bone Healing and Tissue Engineering

**DOI:** 10.1101/2022.11.15.513563

**Authors:** Jacob D. Weston, Brooke Austin, Hunter Levis, Jared Zitnay, Jeffrey A. Weiss, Brandon Lawrence, Robby D. Bowles

## Abstract

Engineered bone tissue synthesized from mesenchymal stem cell progenitors has numerous applications throughout the fields of regenerative medicine and tissue engineering. However, these multipotent cells offer little tissue-building assistance without differentiation direction from environmental cues such as bone morphogenetic proteins (BMPs). Unfortunately, BMP dosing and environmental cues can be difficult to control both *in vitro* and after *in vivo* delivery. Several BMP antagonists are expressed by cells in response to BMP dosing that bind extracellular BMPs and reduce their effective concentration. Here, we use CRISPR-guided gene-modulation technology to downregulate the expression of three BMP antagonists, noggin, gremlin-1, and gremlin-2, in adipose-derived stem cells (ASCs). We show that regulating noggin using this method results in ASC osteogenesis without the need for exogenous growth factors. To demonstrate the versatility and the precision capabilities of these engineered cells, we employ them with CRISPRa multiplex-engineered chondrogenic cells as a proof-of-concept tissue engineering application by creating a tissue gradient similar to the fibrocartilage-to-mineralized-fibrocartilage gradient in the tendon/ligament enthesis or intervertebral disc attachment. In doing so, we show that multiple CRISPR multiplex engineered cell types can be utilized in concert to provide a high degree of tissue developmental control without the use of exogenous growth factors.

## INTRODUCTION

Though bone is renowned for its ability to self-heal with minimal manipulation and sufficient time, 10% of bone fractures do not heal properly^1^. Additionally, some regenerative medicine strategies have struggled to properly control bone mineral deposition^2–5^. 500,000 bone-grafting procedures are performed every year in the United States to encourage bone repair^6^. Using autologous tissue in many of these procedures has long been the gold standard, but this approach can have serious drawbacks, including donor site morbidity, limited supply, and long surgical times^7,8^. Cell therapies utilizing the regenerative properties of multipotent mesenchymal stem cells have also been considered. While stem cells are notorious for their capacity to differentiate into multiple cell types, these cells rely on environmental cues to initiate a commitment to a particular cell fate. Without the differentiation direction often provided by exogenous growth factors or highly specific culture media, stem cell therapies often fail to achieve their desired outcome^9,10^. In recent years, recombinant bone morphogenetic proteins (BMPs) have garnered widespread use in bone healing applications^7,11,12^. BMPs promote bone growth and subsequent healing by recruiting nearby stem cells and driving their osteogenic differentiation^13,14^.

However, this process also results in the upregulation of several BMP antagonists, including gremlin-1 (grem1), gremlin-2 (grem2), and noggin (nog)^15–20^. These antagonists bind to extracellular BMPs, thereby preventing them from interacting with their receptors and reducing their effective concentration^15,16,19,20^. Predictably, this antagonistic action retards osteogenesis. The prevailing clinical approach has been to overwhelm antagonists by applying supraphysiological doses of recombinant BMPs^2,11,21,22^. However, high doses of BMPs have been associated with severe side effects^2,21,22^. Utilizing CRISPR epigenome editing we can interrupt BMP antagonism in order to direct stem cell phenotype toward osteogenesis for bone healing and other applications. A cell therapy that involves the delivery of cells sensitized to endogenous BMPs by reducing BMP antagonism could provide a safer and more effective strategy for promoting bone healing.

Until recently, regulating endogenous gene expression has been accomplished using technologies that have been deservedly criticized for their abundance of off-target effects, which can reduce efficiency and cause unwanted, non-specific interference with other genes^23,24^. Clustered regularly-interspaced short palindromic repeats (CRISPR) technology has emerged in recent years as a highly efficient method for targeting specific loci. CRISPR interference and activation systems utilize a nuclease-deficient variant of the CRISPR-associated protein 9 (Cas9), known as deactivated Cas9 (dCas9). dCas9 is fused to effector molecules that are capable of altering gene expression at the target site. The Krüppel-associated box (KRAB) domain can effectively downregulate target genes when fused as an effector molecule to the dCas9 protein^25^. With an alternative effector molecule such as VPR, similar CRISPR systems can also be used for highly specific gene upregulation^26^. This grants unprecedented control over gene expression and resultant cell phenotype with fewer off-target interactions than the alternatives^27–30^.

For bone healing and tissue engineering, a homogenous population of osteoblasts is desirable to form solid, bridging bone tissue. However, terminally differentiated osteoblasts lack the self-renewal and proliferation abilities that stem cells possess, which makes obtaining a sufficient quantity of them difficult. Adipose-derived stem cells (ASCs) are a type of multipotent stem cell that can be easily and non-invasively obtained from adult adipose tissue. In the early stages of stem cell differentiation, cell fate is determined by the set of genes that are being expressed in response to the local cell environment. By altering specific elements of the transcriptome of ASCs, their differentiation can be guided to achieve a functionally homogenous population of a single, osteoblastic cell type.

Cells with directed osteogenic differentiation would have numerous clinical applications outside of bone healing as well, even in non-homogenous tissues. Such applications would include osteochondral transition zones such as in the attachment of the intervertebral disc (IVD) to the endplate of the vertebral body in the spine, or in tendon and ligament entheses that serve to attach musculoskeletal tissues to bone. These transition zones are comprised of multiple cell types spatially distributed in a gradient that effectively transitions the tissue from a dense, fibrous connective tissue to uncalcified fibrocartilage, to calcified fibrocartilage, and finally to bone^31,32^. We have previously shown that multiplex CRISPR-guided gene activation of aggrecan (ACAN) and collagen type-II (Col2) can effectively increase the deposition of key cartilage ECM components in ASCs^33^. Additionally, our collaborators have created high-density anisotropic collagen scaffolds that mimic natural connective tissue and were shown to be compatible with ASC growth^34^. Co-culture of the ACAN/Col2-upregulated ASCs alongside an osteogenic ASC cell type with the appropriate spatial distribution on such a relevant scaffold would demonstrate the precision engineering capabilities of cells edited with CRISPR-guided gene modulation, and could be a significant step toward tissue-engineering applications.

Here, we demonstrate the potential of CRISPR gene-regulating strategies in targeting BMP antagonists in ASCs. We show that by doing so we are able to effectively regulate these specific genes, and that by downregulating noggin, we were able to robustly affect a phenotypic change that resulted in improved osteogenic differentiation both in monolayer and in a clinically relevant three-dimensional monoculture. As a demonstration of potential tissue engineering applications, we then employ the noggin-knockdown ASCs in co-culture with ACAN/Col2-upregulated ASCs in an IVD attachment or tendon/ligament enthesis tissue-engineering application. These results showcase the capabilities of CRISPR-guided gene modulation technologies for directing the differentiation of multipotent stem cells, and also provide the basis for cell therapies that could be used clinically for bone-healing and tissue engineering applications.

## MATERIALS AND METHODS

### Experimental Overview

Experiments were performed to apply CRISPR epigenome editing to modify the expression of BMP antagonists in ASCs with the purpose of altering cell phenotype to encourage osteogenic differentiation. First, lentiviral CRISPR epigenome editing vectors that targeted the promoter region of the BMP antagonists, gremlin-1 (grem1), gremlin-2 (grem2), and noggin (nog), were designed, built, and tested for the ability to regulate target gene expression in ASCs. The CRISPR-modified ASCs were then cultured in monolayer and in 3D cultures to demonstrate improved osteogenesis by evaluating alkaline phosphatase (ALP) activity, calcium deposition, and microcomputed tomography (micro-CT) imaging.

### Lentiviral CRISPR Epigenome-Editing Vector Construction

Lentiviral CRISPR epigenome-editing vectors targeting the nog, grem1, and grem2 gene promoter regions, and a non-target control (NTC), were designed and built using a previously described method^35,36^. Briefly, target-specific protospacers were cloned into the hUBC-dCas9-KRAB-T2A-GFP vector under the control of an hU6 promoter (Figure 1A). The protospacer sequences were designed using GT-Scan based on anticipated off-target interactions and coincidence with DNase-I hypersensitive regions as determined by the UCSC genome browser at the promoter region of each target gene (Figure 1B)^37–39^. Plasmids were amplified in NEB Stable competent E. coli and purified using a QiaPrep Spin Miniprep kit (Qiagen, Venlo, NL). All plasmids were subjected to Sanger sequencing to verify the presence of the correct sing guide RNA (sgRNA) sequences.

**Figure 1:**
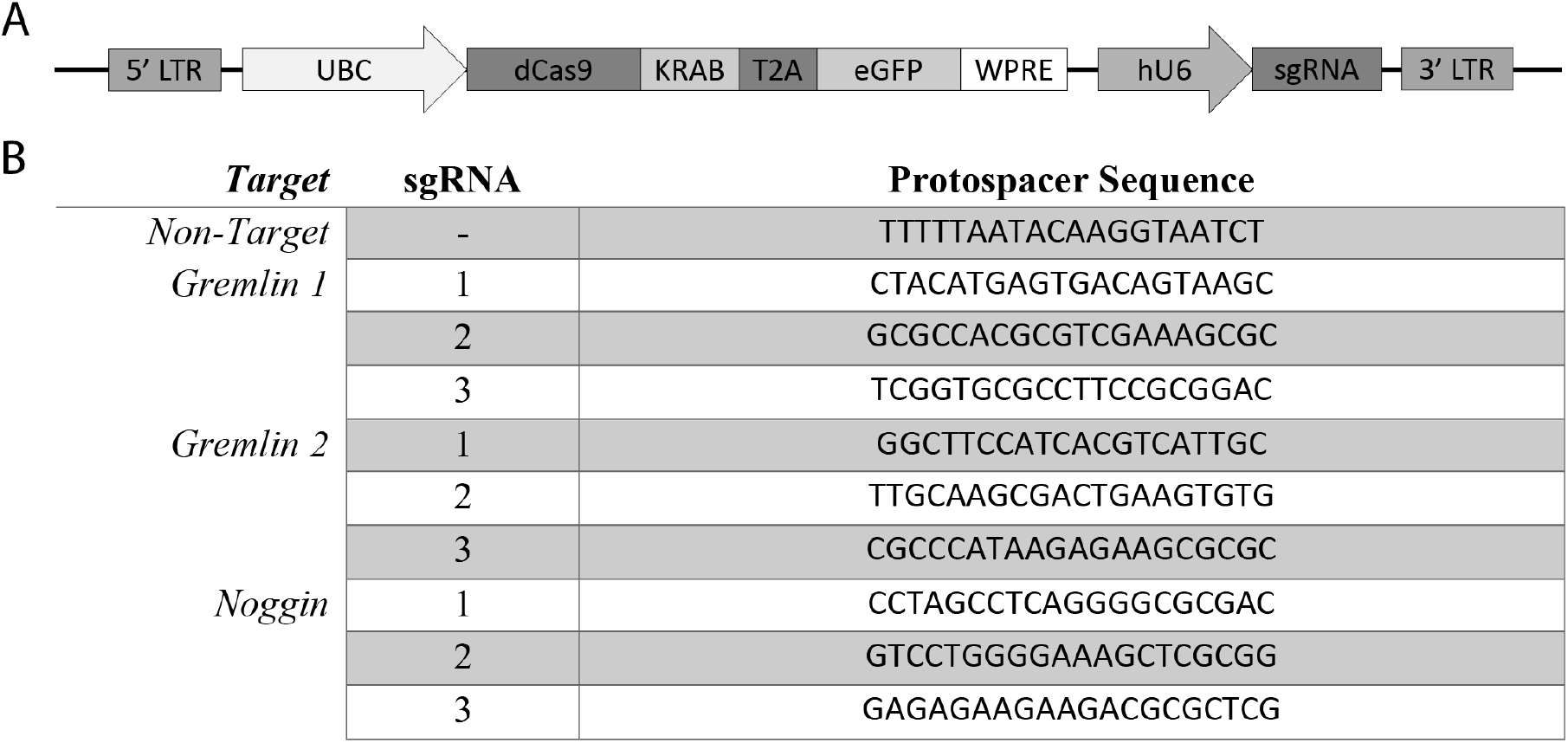
(A) CRISPR epigenome editing lentiviral expression cassette. (B) sgRNA sequences for each of the BMP-antagonist target genes.

### Lentivirus Production

Lentiviral vectors co-expressing dCas9-KRAB-T2A-eGFP and targeting sgRNAs were produced as previously described^36^. Briefly, HEK293T cells were seeded at 62,500 cells/cm2 and cultured in high-glucose DMEM (Gibco) supplemented with 10% FBS (Hyclone, Logan, UT). The following day, they were co-transfected with the lentiviral CRISPR epigenome-editing expression plasmid, psPAX2(Addgene, plasmid #12260), and pMD2.G(Addgene, plasmid #12259). Lentiviral supernatant was collected, concentrated with Lenti-X concentrator (Clontech, Mountain View, CA), and stored at −80C until use.

### Creation of Epigenome-Edited ASC Cell Lines

Immortalized ASCs (SCRC-4000; ATCC, Manassas, VA) were seeded (10,000 cells/cm^2^) onto BioLite cell-culture plates (Thermo Scientific, Waltham, MA) and cultured in complete ASC growth medium (ATCC) for 24 hours (37C, 5% CO2). 125 ul of concentrated virus, supplemented with 4ug/ml polybrene (Sigma-Aldrich, St. Louis, MO), was added to the cell culture media and incubated for 24 hours. Cells were then washed 3x with PBS and cultured for one week prior to sorting. Transduced cells were subjected to fluorescence-activated cell sorting for eGFP fluorescence to obtain homogenous populations of epigenome-edited cells.

### qRT-PCR

Gene expression in the epigenome-edited cells (n = 3) was evaluated using quantitative reverse transcriptase (qRT) PCR. Following a one-week culture period, cells were lysed and total RNA was extracted from cell lysates using a Purelink RNA micro kit (Invitrogen). cDNA was synthesized from the total RNA using a High-Capacity cDNA Reverse Transcription Kit (Applied Biosystems, Foster City, CA), and then used in the qRT-PCR reaction. Target gene expression was determined using TaqMan Gene Expression Assay probes (Applied Biosystems), normalized to GAPDH expression, and analyzed using the comparative Ct (ΔΔCt) method to produce fold-change values relative to the NTC group.

### ALP Activity Quantification and Staining

#### Monolayer Cell Culture of epigenome-edited ASCs

Epigenome-edited cell lines showing the greatest downregulation of nog, grem1, or grem2, were seeded in 96-well plates (5,000 cells/cm^2^) in growth medium. Once the cells reached confluency, recombinant BMP-2 (R&D Systems, Minneapolis, MN) at varying doses (0, 30, or 100 ng/ml) was added to the appropriate wells. Cells were cultured for seven days, with fresh BMP-2 addition and media changes every 2-3 days.

#### Quantitative ALP Activity Assay

After seven days of culture, the sorted cells were lysed (0.2% Triton X-100; Acros Organics, Waltham, MA), and ALP activity was determined using a commercially available assay (QuantiChrom ALP Assay Kit; BioAssay Systems, Hayward, CA) according to the manufacturer’s protocol (n = 6).

#### ALP Staining

Following 7 days of culture, cells were rinsed with PBS and then fixed with neutral-buffered formalin for 60 seconds (n = 2). Cells were then washed with 0.05% Tween 20 in PBS. The fixed cells were incubated with SIGMAFAST ALP substrate solution (Sigma) for 10 minutes at room temperature. The substrate solution was discarded and the cells were washed again with 0.05% Tween 20. PBS was then added to each well before imaging. Images were captured with a GelCam 315 Camera (Analytik Jena US, Upland, CA).

### Monolayer Alizarin Red Staining and Image Analysis

Sorted cells were cultured for 21 days following initial BMP-2 dosing (n = 4). The cells were then rinsed (PBS) and fixed (10% neutral-buffered formalin, 30 minutes at room temperature). After fixation, cells were rinsed (PBS) and incubated in Alizarin Red solution (20g/L in water, pH 4.2, 45 minutes at room temperature). Following this incubation, the cells were rinsed 4x with water and then imaged in PBS as described in the ALP staining section.

Images were processed and analyzed using ImageJ software^40^. Greyscale images were first inverted, and then background subtraction was applied using the rolling ball method with a radius of 12.5 pixels^41^. Representative portions of the wells were selected, and the blank-subtracted mean grey values were quantified for each well to produce the reported values with arbitrary intensity units.

### Osteogenic Differentiation of Noggin-targeted Cells in 3D Culture

Sterile type-I bovine collagen sponges (Integra LifeSciences, Plainsboro, NJ) were aseptically cut into 2.5mm cubes. These cubes (n=4) were submerged in a cell suspensions at 10,000,000 cells/ml. The suspension with the sponges was mixed, and the loaded sponges were transferred to individual wells of a V-bottom plate. The plate was then incubated at 37C for 30 minutes before transferring them to wells containing growth medium. After one week of culture, half of the samples were changed to osteogenic medium (low-glucose DMEM, 10% MSC FBS, 2mM GlutaMax, 100U/ml penicillin, 0.1 mg/ml streptomycin, 10nM dexamethasone, 50μM 2-phospho-L-ascorbic acid, 10mM β-glycerophosphate, and 33μM phenol red). The cells were cultured for another 21 days in these conditions, after which they were fixed with 10% neutral-buffered formalin for 24 hours at room temperature and then submerged in 70% ethanol prior to histological and microcomputed tomography analysis.

### Osteogenic Differentiation on Anisotropic High-Density Collagen Gels

#### Synthesis of Anisotropic High-Density Collagen Gels

Anisotropic high-density type-I collagen gels were created by means of biaxial compression as previously described^34^. Briefly, the gels are made in a three-phase process. First, a solution containing acid-solubilized type-I collagen (6 mg/ml) and porogens made of a polyurethane elastomer is polymerized in a rectangular mold. Second, the gels are serially compressed along two orthogonal directions by means of a mechanical compression apparatus to densify the material and align the collagen fibrils. Finally, the gels are rinsed three times in 80% ethanol for 24 hours at 5 °C to remove the porogens. Prior to cell seeding, the gels were rinsed twice in PBS at 37 °C for 24 hours, followed by a single rinse in cell culture medium at 37 °C for 24 hours.

#### Cell Seeding and Culture on High-Density Collagen Gel Cubes

Cell suspensions of non-target and noggin-targeted cells at 10,000,000 cells/mL were made for each cell type. The high-density collagen gels were cut into 2.5 mm cubes using sterile shears. The cubes (n=4) were submerged in their appropriate cell suspension and cells were allowed to attach to the cubes for 1 hour at 37C, during which time they were briefly agitated every 20 minutes to encourage uniform seeding on all sides of each gel. The gels were then transferred to individual wells and cultured at 37C in growth medium for one week. The medium was then replaced with osteogenic medium cultured for 6 weeks with media changes every 2-3 days.

#### Gradient Cell Seeding on Full-Size Collagen Gels

A cell-seeding apparatus was designed using SolidWorks software, and 3D printed at high-resolution on a Formlabs Form 2 3D printer (Formlabs, Somerville, MA). The apparatus featured 5 cell-seeding channels measuring 3×12×5 mm each. For cell seeding, the apparatus was placed directly on the 15 mm x 20 mm collagen gel and a weight was placed on top of it to ensure a proper seal between the apparatus and the gel. 5 different cell suspensions were made, all at 10,000,000 cells/mL. These suspensions were comprised of noggin-knockdown cells and ACAN/Col2-upregulated cells in the following ratios: 1:0, 3:1, 1:1, 1:3, and 0:1. 150 μL of each cell suspension was added to its respective channel in the cell-seeding apparatus, and the cells were incubated at room temperature for 30 minutes to allow for cell attachment. The cell suspensions were then aspirated and the apparatus was removed from the gel. The cell-seeding process was then repeated on the other side of the gel. After both sides were seeded, the gel was incubated at 37C in growth medium for 48 hours before being cut longitudinally into 3×20 mm strips, perpendicular to the orientation of the cell-seeding channels. The cell-laden strips were cultured in growth medium for 7 days and then in osteogenic medium for 6 weeks, after which they were fixed in 4% paraformaldehyde in PBS for 24 hours.

### Histology

Fixed samples were embedded in paraffin, cut into 7μm (4 μm for high-density gels) sections, and mounted onto slides. Hematoxylin and eosin (H&E), Von Kossa, alcian blue, and alizarin red staining were then performed using established staining protocols^42^. Slides were imaged by bright-field microscopy at 40x magnification using a Nikon Eclipse E400 Microscope (Nikon Inc., Melville, NY) and an Olympus UC50 camera (Olympus Corp., Tokyo, Japan).

Semi-quantitative analysis of Von Kossa- and alizarin red-stained histological sections was performed using ImageJ software. Images were converted to 8-bit greyscale images and a threshold was applied to eliminate background and reduce noise. A representative area of the section was then selected and mean grey values corresponding to staining levels were measured.

### Microcomputed Tomography

The formalin-fixed paraffin-embedded sponges and high-density gel strips were visualized by micro-CT using a Siemens Inveon Preclinical Micro-CT scanner (Siemens AG, Munich, Germany) with the following settings for the sponges and gels respectively: voxel size: 19.5μm, 34.5 μm; voltage: 70kV, 70kV; current: 200μA, 350 μA; exposure: 2500ms, 1200ms.

Volumetric renderings were generated using RadiAnt DICOM Viewer (Medixant, Poznan, Poland). Data were quantified using ImageJ software by measuring the average pixel intensity for each cross-section along the length of each construct. Due to differing conditions at each gel end compared to the rest of the gel, and to better display the properties of the bulk material, the first and last 10% of each high-density gel strip was excluded from quantification. Images were calibrated by scanning a water phantom with the same instrument settings, and raw attenuation coefficients were converted to Hounsfield units by applying a linear transformation based on the readings of air and water from the phantom.

### Statistics

All plots and statistical analyses were generated using ggplot2 (Wickham, 2009) and the base stats packages of R (R Core Team, 2017). For qRT-PCR datasets, statistical significance was determined using one-way or two-way analysis of variance (ANOVA) as appropriate with a Tukey HSD post hoc test. Alpha was set at 0.05 for all statistical analyses.

## RESULTS

### CRISPR Epigenome Editing of BMP antagonist promoter regions in ASCs robustly downregulates expression of the target genes

To identify an effective sgRNA, 3 sgRNAs were designed for the promoter region of each gene of interest. Transduction of ASCs with these vectors demonstrated robust downregulation of the target gene expression at the RNA level (Figure 2), with the most effective guides providing maximum downregulation of 88% (±0.92%, n=3), 58% (±3.9%, n=3), and 92% (±0.65%, n=3) for grem1, grem2, and nog respectively. The cells containing the best-performing sgRNA for each target gene were selected from these data and were used for all downstream differentiation experiments.

**Figure 2:**
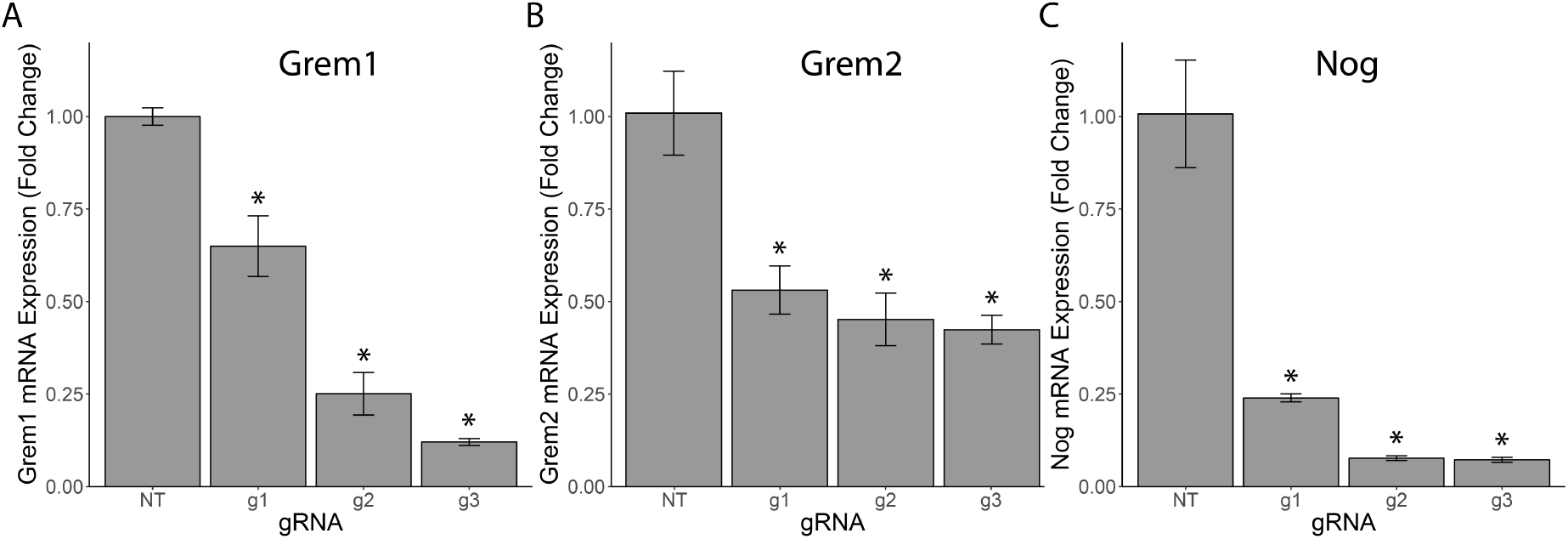
Target gene mRNA expression for each sgRNA normalized to the non-target (NT) group for gremlin 1-(A), gremlin 2-(B), and noggin-targeted (C) cells. * = p<0.05 as compared to the NT group.

### Epigenome Editing at the Noggin Promoter Increases ALP Activity and Calcium Deposition in Monolayer Cultures

The quantitative ALP activity assay revealed that nog-targeted cells demonstrated significantly higher (p<0.05) ALP activity than controls, regardless of the level of exogenous BMP-2 dosing (Figure 3A). ALP levels increased 92, 60, 85, and 53% for 0, 10, 30, and 100 ng/ml BMP-2 doses respectively. For the other antagonist-targeted groups, no increase in ALP activity was observed. Instead, grem1-targeted cells were shown to have significantly decreased ALP activity (p < 0.05) for each of the four BMP-2 doses, and grem2-targeted cells also generally had lower ALP activity, with the 10 and 100 ng/ml BMP-2 groups being significantly reduced (p<0.05). Qualitative analysis of ALP staining further confirmed the results of the quantitative assay (Figure 3B), showing general trends of darker staining in the noggin-targeted group, lighter staining in the grem1 group, and little difference in the grem2 group, as compared to controls.

**Figure 3:**
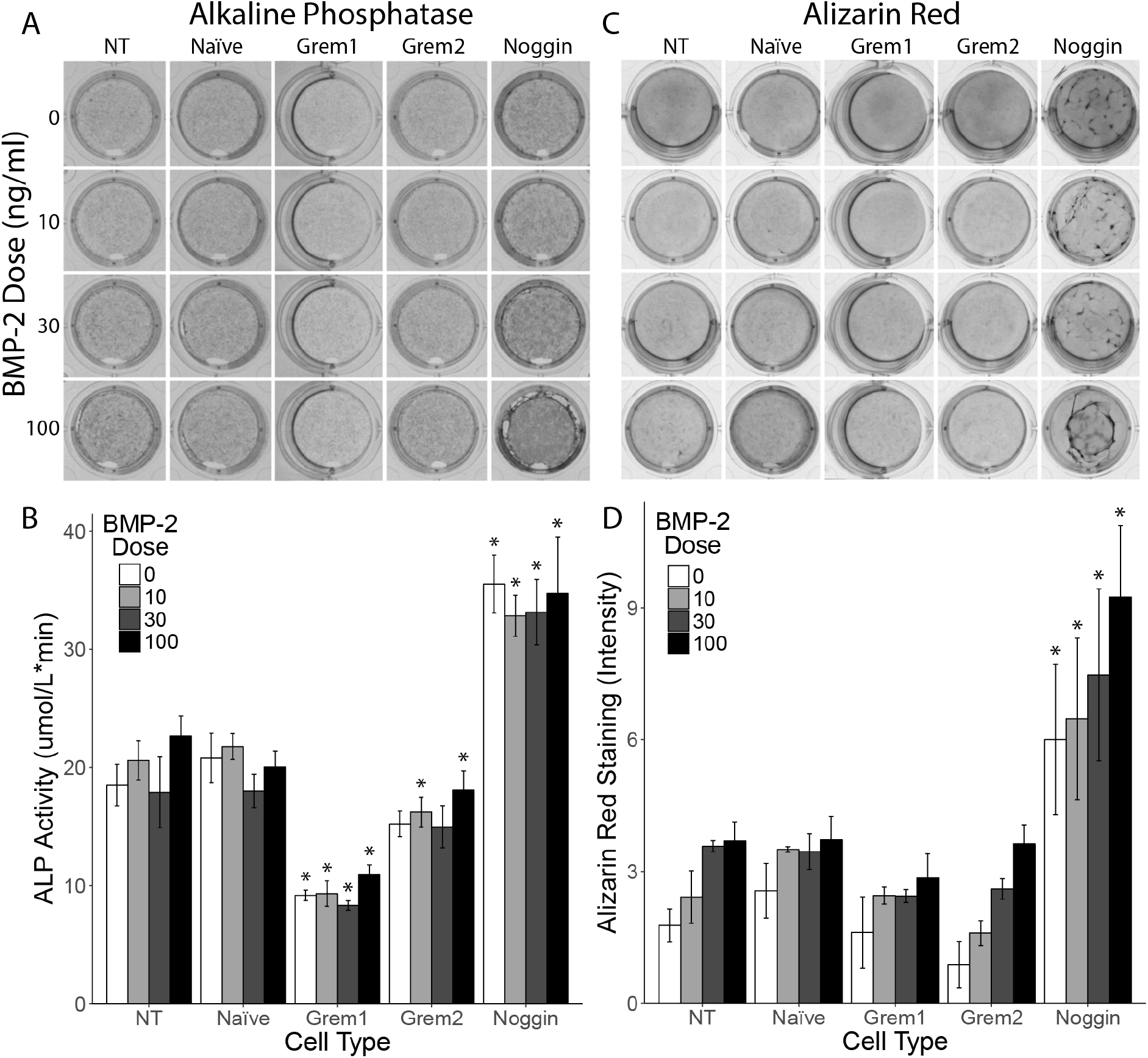
(A) ALP activity staining and (B) quantitative assay for cells cultured with varying doses of BMP-2. (C) Alizarin red staining of each cell line and (D) semi-quantitative image analysis of alizarin red staining for calcium deposition. * = p<0.05 as compared to the NT group with the equivalent BMP-2 dose.

Noggin-targeted cells demonstrated markedly increased calcium deposition and mineralized nodule formation as compared to NTCs, while all other groups showed little to no difference (Figure 3C). Semi-quantitative analysis of the images showed that these increases in calcium deposition were statistically significant (p < 0.001; Figure 3D). Increases in calcium deposition in the noggin-targeted cells were calculated to be 197, 134, 109, and 170% for 0, 10, 30, and 100 ng/ml BMP-2 doses respectively, relative to the NTC.

### Epigenome Editing at the Noggin Promoter Increases Mineralization in 3D Cultures

H&E staining of 3D culture cross sections showed little difference between the NTC and the noggin-targeted cells in either growth or osteogenic media. Though all samples started approximately the same size, the cultures grown in osteogenic medium for both cell types were visibly smaller at day 21 than their counterparts cultured in growth medium (Figure 4).

**Figure 4:**
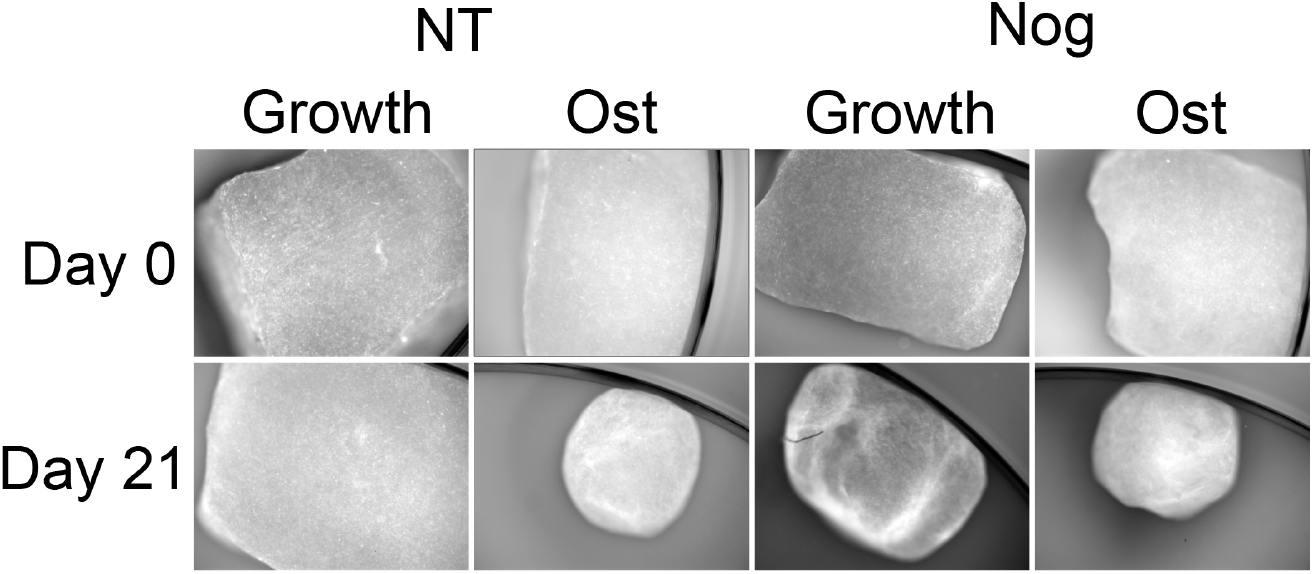
Photomicrographs at 40x magnification of the collagen sponges seeded with non-target (NT) and noggin-targeted (Nog) cells at day 0 and day 21 of osteogenic induction.

Von Kossa staining revealed a dramatic difference in mineralization between non-target and noggin-targeted cells (Figure 5A). Sponges seeded with noggin-targeted cells grown in osteogenic medium had extensive mineralization throughout the sponge at day 21, which was entirely absent in the non-target group. This marked increase in mineralization was further confirmed by alizarin red staining (Figure 5A). A statistically significant (p < 0.05) difference between the noggin-targeted cells cultured in osteogenic medium and all other treatment groups was seen from the semi-quantitative analysis of the images of the Von Kossa- and alizarin red-stained histological sections (Figure 5B). Furthermore, this analysis also showed a significant increase in calcium deposition in the noggin-targeted group cultured in growth medium, indicating an increase in osteogenic potential even in the absence of other osteogenic factors.

**Figure 5:**
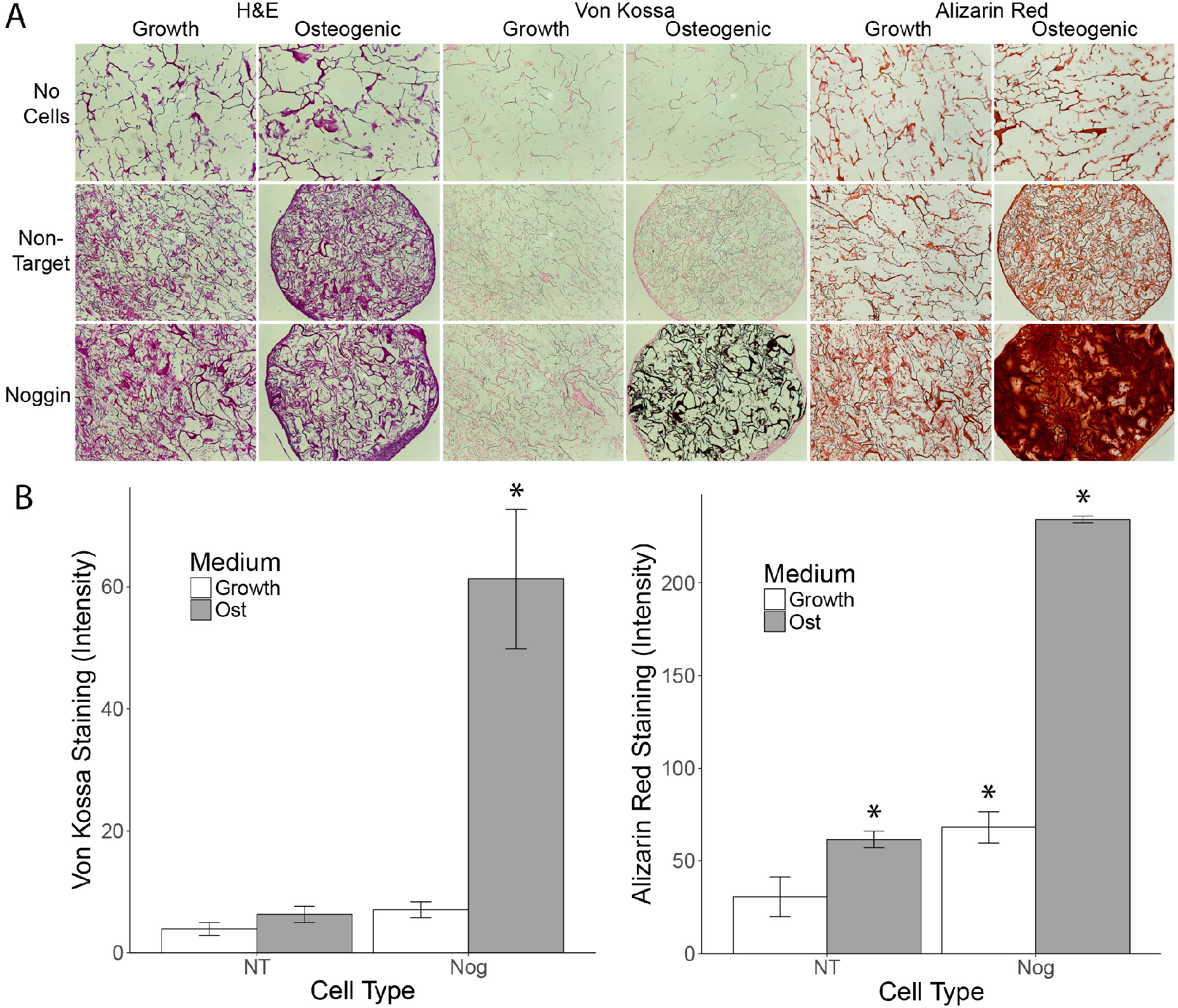
(A) H&E-, Von Kossa-, and alizarin red-stained histological sections of the fixed collagen sponges after 21 days of culture in either growth or osteogenic medium. (B) Semi-quantitative image analysis of Von Kossa-stained sections. (C) Semi-quantitative image analysis of alizarin red-stained sections. (*= p<0.05 as compared to the NT-growth medium group)

Micro-CT imaging revealed dramatic qualitative and quantitative differences between the noggin-targeted and the NTC groups cultured in osteogenic medium (Figure 6A-B). Noggin-targeted sponges in osteogenic medium had a significantly higher (p < 0.001) average radiographic density of 101.7±62.5 HU compared to −398.5±21.9 HU in the NTC group. Both of the groups cultured in standard growth medium also had low densities of −403.1±7.1 HU and - 412.1±3.9 HU for the noggin-targeted and NTC groups respectively. No statistically significant difference was observed between these values.

**Figure 6:**
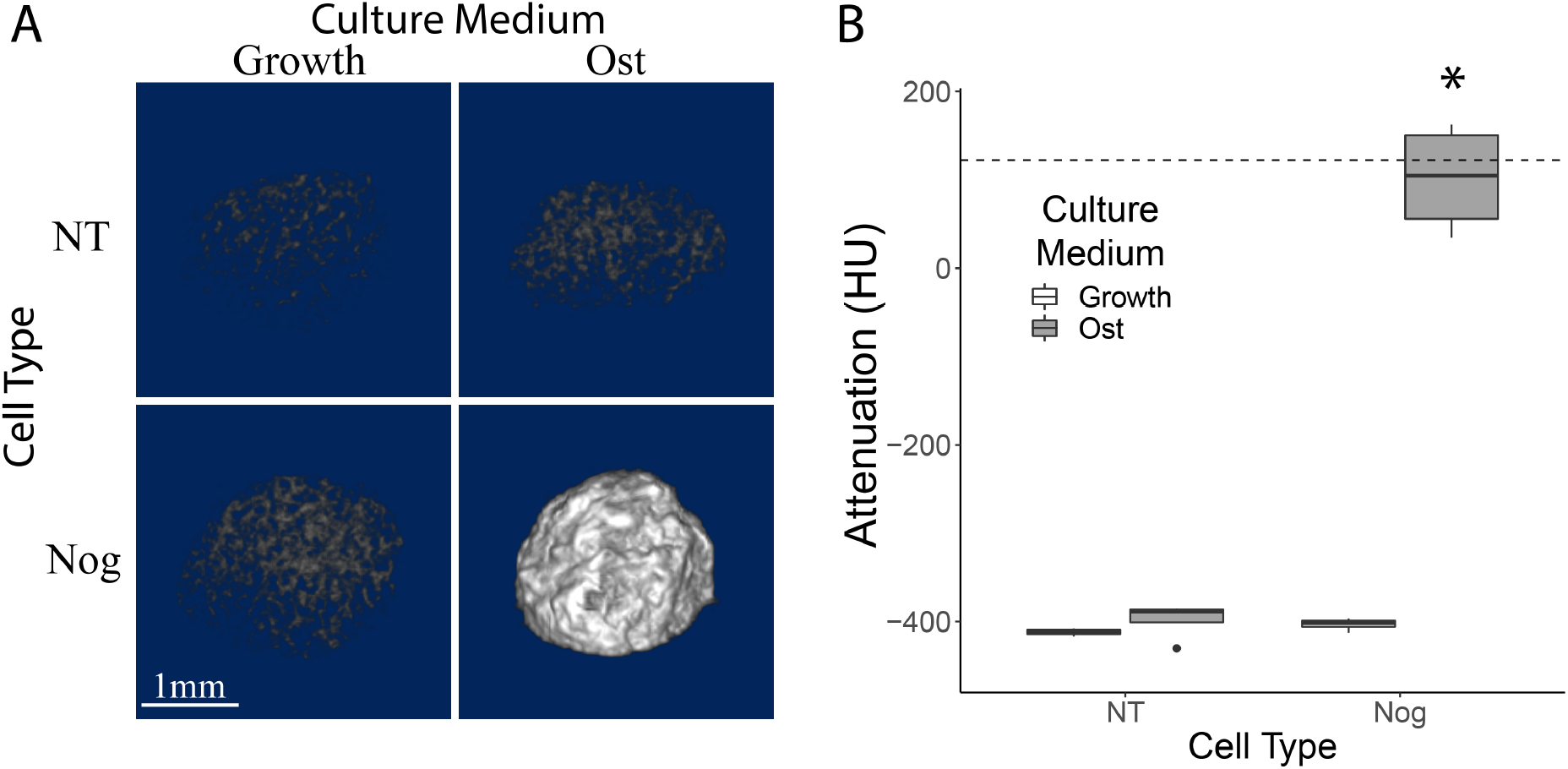
(A) Volumetric renderings of micro-CT scans of noggin-targeted and non-target cells on collagen sponges after 21 days of culture in osteogenic medium. (B) Radiographic density of each of the treatment groups as measured by micro-CT. * denotes a statistically significant difference (p < 0.001) as compared to each of the other groups. The dashed line is the average radiographic density of human vertebral cancellous bone (121HU) from the literature^43^.

### Gradient on Anisotropic Collagen Scaffolds

In single culture, the noggin knockdown cells and ACAN/Col2 cells showed increased calcium and sulfated glycosaminoglycan (sGAG) deposition respectively on the high-density gels (Figure 7A-B). For the co-culture experiments, the cell-seeding apparatus (Figure 7C) allowed the two cell types to be seeded in a gradient across the gel (Figure 7D-E).

**Figure 7:**
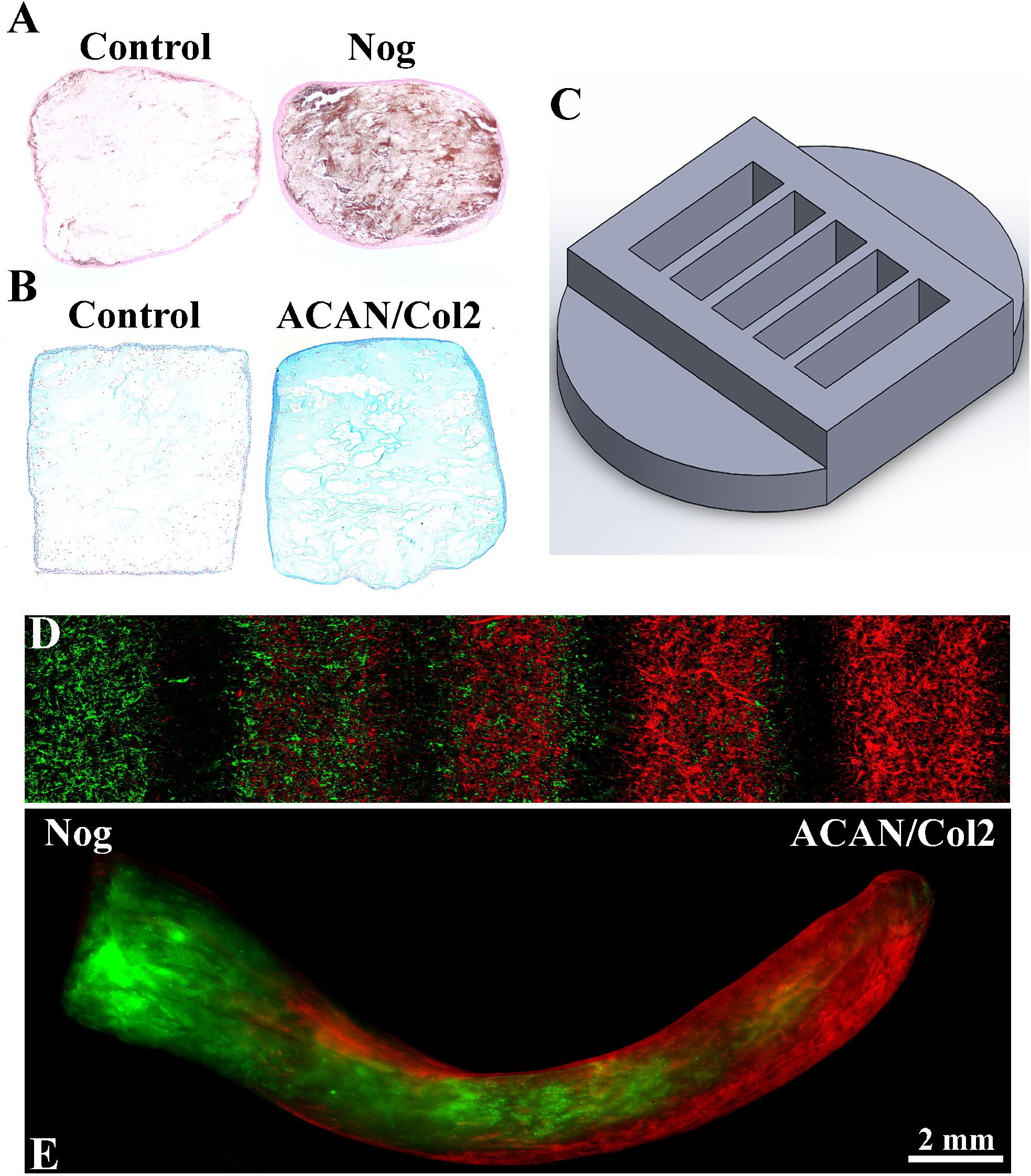
(A) Von Kossa and (B) alcian blue staining of histological sections of high-Density anisotropic collagen gel cubes after 6 weeks. (C) The device used for seeding the cells in a gradient. Noggin-knockdown [green] and ACAN/Col2-upregulated [red] cells 24 hours (D) and 2 weeks (E) after seeding on the full-size collagen gel.

Quantified micro CT scans of the anisotropic collagen gels seeded with cell gradients revealed a density gradient that correlated with noggin-knockdown cell seeding (Figure 8A-B). Optimal sections of the gels (n=3), based on the density gradient, had a density of 401.8±47.3 HU on one end and 244.1±60.4 HU on the other with a density slope of 16.4±4.3 HU/mm over an average length of 10.2±3.7 mm. Von Kossa staining showed substantial mineralization on the end seeded with noggin-knockdown cells and gradually decreasing mineralization toward the other end (Figure 8C). However, the mineralization was mostly confined to the external surfaces of the gels, thereby limiting the utility of these histological stains. Alcian blue staining of the sections revealed that sGAG deposition was distributed throughout the gel, though no gradient was observed (Figure 8D).

**Figure 8:**
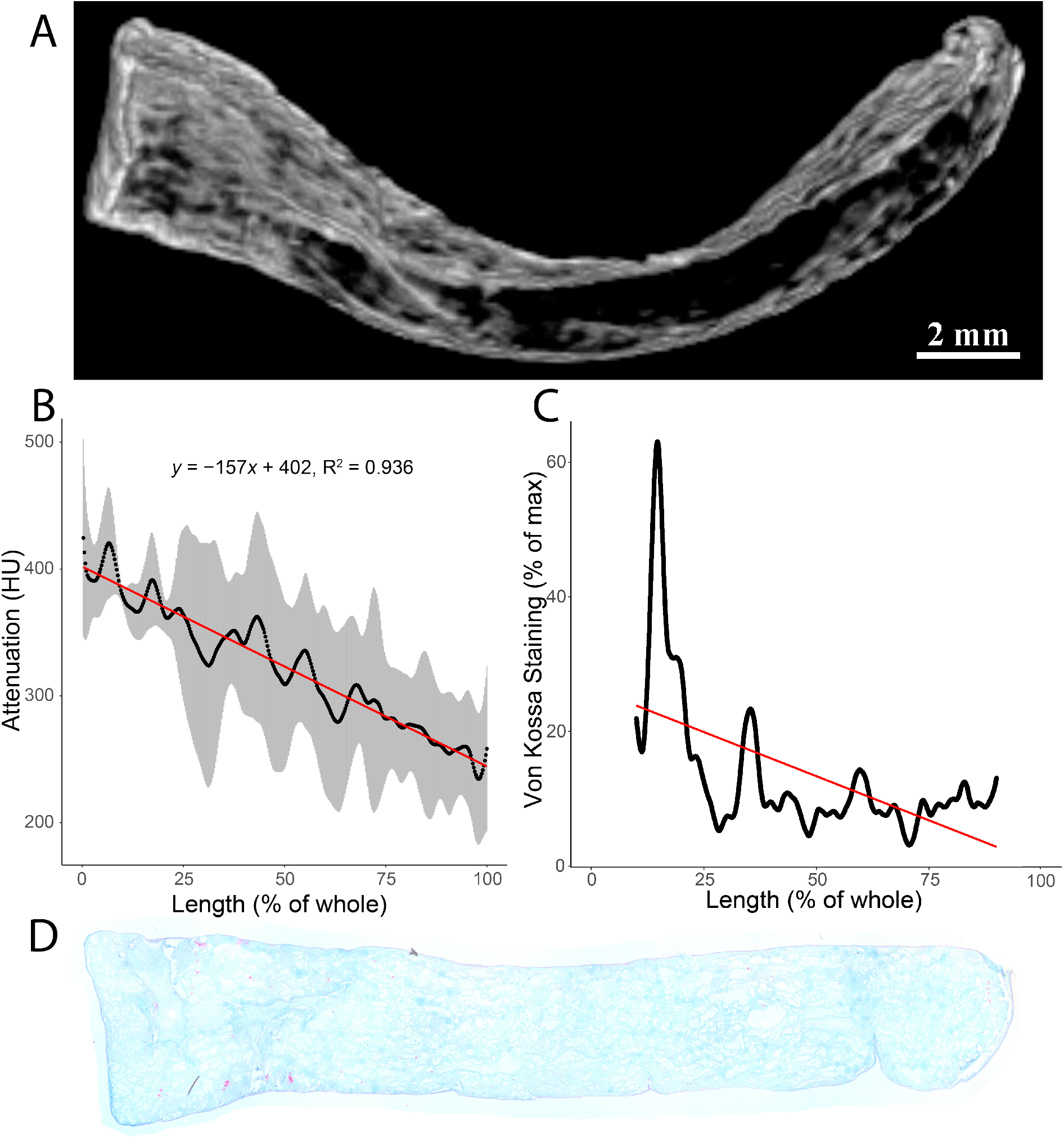
(A) Volumetric rendering of one of the gels showing the regions of the gel with the highest densities. (B) Scatter plot of the quantified micro-ct scans showing the signal attenuation of the portions of the gels with the best density gradient. Shaded region shows standard deviation and the red line shows the linear trendline with the inset equation. (C) Semi-quantitative analysis of the von kossa staining of a histological section from one of the gels. (D) Alcian Blue staining after 6 weeks of culture.

## DISCUSSION

Despite the enormous potential of multipotent stem cells in therapeutic and tissue engineering applications, these cells are of little use unless their differentiation can be controlled. Thus far, such control over their differentiation has largely been accomplished by dosing with exogenous growth factors. While these methods have been effective in some cases, there are many instances where such approaches are not practical. The results shown here demonstrate effective control over osteogenic differentiation of a multipotent stem cell line without exogenous growth factors using CRISPR-guided gene modulation, as well as a proof-of-concept application where the engineered cells are utilized to create a tissue gradient of calcified to uncalcified fibrocartilage.

We first reported improved ASC osteogenesis with CRISPRi knockdown of noggin at the 2019 Orthopaedic Research Society Annual Meeting^44^. Since that time, others have shown osteogenic improvement in rat ASCs targeting noggin using similar CRISPRi systems but also delivering vectors overexpressing BMP-2^45^. Remarkably, with the high degree of noggin knockdown in our system, the cells exhibited a marked improvement in osteogenic phenotype, with increased ALP activity and calcium deposition even without the additional growth factors that others have needed^45^. This suggests that by knocking down noggin expression to this degree, even endogenous levels of BMPs and other osteogenic factors become capable of driving osteogenesis in these cells. The osteogenic capacity of these modified cells was further demonstrated in clinically relevant three-dimensional cultures. Resorbable type-I collagen sponges have been used in the clinic in spinal fusion procedures for decades, serving as scaffolding for bone-forming cells and also as carriers for retaining osteogenic growth factors at the treatment site. Here, we have shown dramatic improvements in osteogenic differenation on these same scaffolds, again without the addition of any exogenous osteogenic growth factors, with mineralization levels approaching those seen in natural bone. In the literature, the radiographic density of healthy bone tissues varies, with normal human vertebral cancellous bone having been reported to have an average radiographic density of 121±42 HU ^43^, while other types of cancellous bone have densities around 300 HU ^46,47^. When grown on the collagen sponges, the noggin-targeted group cultured in osteogenic medium was the only group to have any values of such magnitude, with 40.8±8.3% of the total sample volume having a radiographic density of 100 HU or greater, and 21.3±7.2% of the volume having a density of 300 HU or greater.

In culturing the cells in a gradient on the high-density collagen gels, we not only showed a potential tissue-engineering application for these modified cells on a relevant scaffold, but also that the noggin-targeted cells were capable of osteogenic differentiation in co-culture with a non-osteogenic cell type. As has been previously shown in chondrogenic pellet cultures^33^, the ACAN/Col2-upregulated cells demonstrated improved deposition of sGAGs on the gels. However, despite being seeded in a gradient across the gels like the noggin-targeted cells, we did not see a gradient in alcian blue staining like we saw with density and mineralization. While this was surprising, it wasn’t entirely unexpected. The literature has shown the difficulties of proteoglycan retention in type I collagen scaffolds, especially in the absence of additives included specifically to anchor them in place^48,49^. With much of the cell growth localized on the surface of the gels and nothing anchoring them in place, sGAGs likely diffused throughout the gel and into the culture supernatant instead of staying fixed in place. Despite the lack of a gradient of proteoglycans, their distributed presence combined with the calcification gradient on these gels still serves to mimic the fibrocartilage to mineralized fibrocartilage region of osteochondral transition zones. Future work will include the exploration of different cell seeding methods, gel porosities, and further CRISPR-guided gene modulation to improve cell distribution and proteoglycan retention throughout the gels.

In summary, we have shown that CRISPR-guided gene modulation is an effective method for driving osteogenic differentiation in ASCs without the use of exogenous growth factors. We have also employed these pro-osteogenic cells in two different 3D cultures that have direct clinical applications in bone healing and tissue engineering, thus demonstrating their versatility and osteogenic potency. While much work remains to be done, these types of systems have clearly shown that they can provide a means for developing effective cell therapies for the treatment of problematic orthopedic conditions.

## Acknowledgements

Funding for this research was provided by the University of Utah Department of Orthopaedics and the Lumbar Spine Research Society.

Research reported in this publication utilized the Biorepository and Molecular Pathology Shared Resource at Huntsman Cancer Institute at the University of Utah and was supported by the National Cancer Institute of the National Institutes of Health under Award Number P30CA042014. The content is solely the responsibility of the authors and does not necessarily represent the official views of the NIH.

